# Molecularly Distinct Innexin Gap Junction Channels and Undocked Hemichannels Regulate Glia Morphology and Function in *Caenorhabditis elegans*

**DOI:** 10.64898/2026.09.21.753071

**Authors:** Arunima Sen, Marlyn Xavier Mascarenhas, Sushrita Roy, Ananya Bandyopadhyay, Abhishek Bhattacharya

**Affiliations:** National Centre for Biological Sciences - TIFR, Bangalore, India

**Author notes:** Correspondence to: Abhishek Bhattacharya.

## Abstract

Glial cells across species orchestrate nervous system development, maintenance, and function through ionic and metabolic crosstalk. These cells express multiple gap junction and hemichannel components. However, the mechanisms by which glia communicate utilizing these channel components to support nervous system architecture and function remain poorly understood. By studying GLR glial cells in *Caenorhabditis elegans*, we show that individual glial cells employ distinct innexin channel configurations to support specific functional roles, utilizing discrete downstream cellular mechanisms. We find that while innexins UNC-7and UNC-9 function as gap junction channels, the innexin INX-18 operates independently to form undocked hemichannels in discrete domains within the same cells. In this combinatorial configuration, UNC-7/UNC-9 gap junctions specifically regulate Synaptobrevin/SNB-1 localization in RME neurons, as previously reported. However, both channel types function non-redundantly to maintain glial morphology by regulating the Calpain/CLP-4 and CDK-5 pathway through cell-autonomous modulation of intracellular calcium levels. In contrast, INX-18 hemichannels in GLR glia regulate high-salt-induced paralysis behavior through distinct downstream cellular mechanisms, potentially in conjunction with tyramine signaling. Altogether, our findings demonstrate a broader repertoire of innexin channel utilization at the individual glial cell level, supporting specific aspects of nervous system maintenance and functioning.

## INTRODUCTION

Coordinated nervous system function relies on precisely regulated intracellular and intercellular communication networks. Across species, intercellular crosstalk between neurons and glial cells regulates nervous system development, maintains ionic homeostasis and buffering conditions necessary for accurate information flow^1–6^, as well as preserves overall nervous system architecture^7–11^. Gap junction channels, which are composed of innexins in invertebrates and connexins in vertebrates, serve as a major conduit for glia-mediated coupling. These channels are specialized intercellular connections that enable direct cytoplasmic communication between partnering cells, allowing near-instantaneous ionic and metabolic coupling. Glial cells, including astrocytes and oligodendrocytes across species, also mediate neuro-glial and glia-glia coupling through these channels^12–14^. In *C. elegans*, glial gap junctions have been shown to regulate axon specification in partnering neurons^9^. In addition to gap junction channels, undocked hemichannels formed by connexins and pannexins in vertebrates, and by innexins in invertebrates, facilitate communication between the cytosol and the extracellular milieu^15^. In glial cells, these hemichannels regulate ionic buffering and metabolite homeostasis^15–18^. Pannexin1 hemichannels in astrocytes are implicated in ATP and Ca^2+^ homeostasis^16,17,19,20^ and modulate cell signaling pathways. Disruptions of these channels are associated with neuroinflammation, seizures, epilepsy, and other neurological disorders^21,22^. Moreover, Pannexin1 hemichannels have also been shown to functionally interact with ligand-gated ion channels, such as N-methyl-D-aspartate receptors (NMDARs), and mutually potentiate each other under both physiological and disease conditions^16,23,24^. Interestingly, glial cells across species express multiple connexin, pannexin, or innexin genes simultaneously^6,25,26^. However, understanding how multiple channel components are assembled to form gap junctions and undocked hemichannels that support multifaceted glial functions remains largely unexplored.

The nervous system of *C. elegans* comprises of 56 glial cells, including six GLR glial cells that are mesoderm-derived and, based on gene expression profiles, exhibit characteristics of both astrocytes and endothelial cells. GLR cells sit just posterior to the nerve ring, the main neuropil in the head of the animal. Each of the six GLR glial cells extends a flat, non-overlapping leaf-like process underneath the nerve ring, which makes contact with multiple cell types **(Figure 1A)**. These leaf-like processes encircle the pseudocoelom, thereby separating the nerve ring from the circulatory system of *C. elegans* and establishing a primitive blood-brain barrier-like structure. GLR cells also extend thin, neurite-like long processes from the leaf-like region towards the nose of the animal, which fasciculate with multiple sensory neuron dendrites **(Figure 1A)**^27^. GLR glial cells have been shown to form gap junctions with neurons and muscles. Specifically, gap junction coupling between GLR glia and RME neurons is necessary for proper axon specification by maintaining the cytoskeletal organization in these neurons through regulating calcium signaling and calcium-dependent microtubular dynamics.

**Figure 1:**
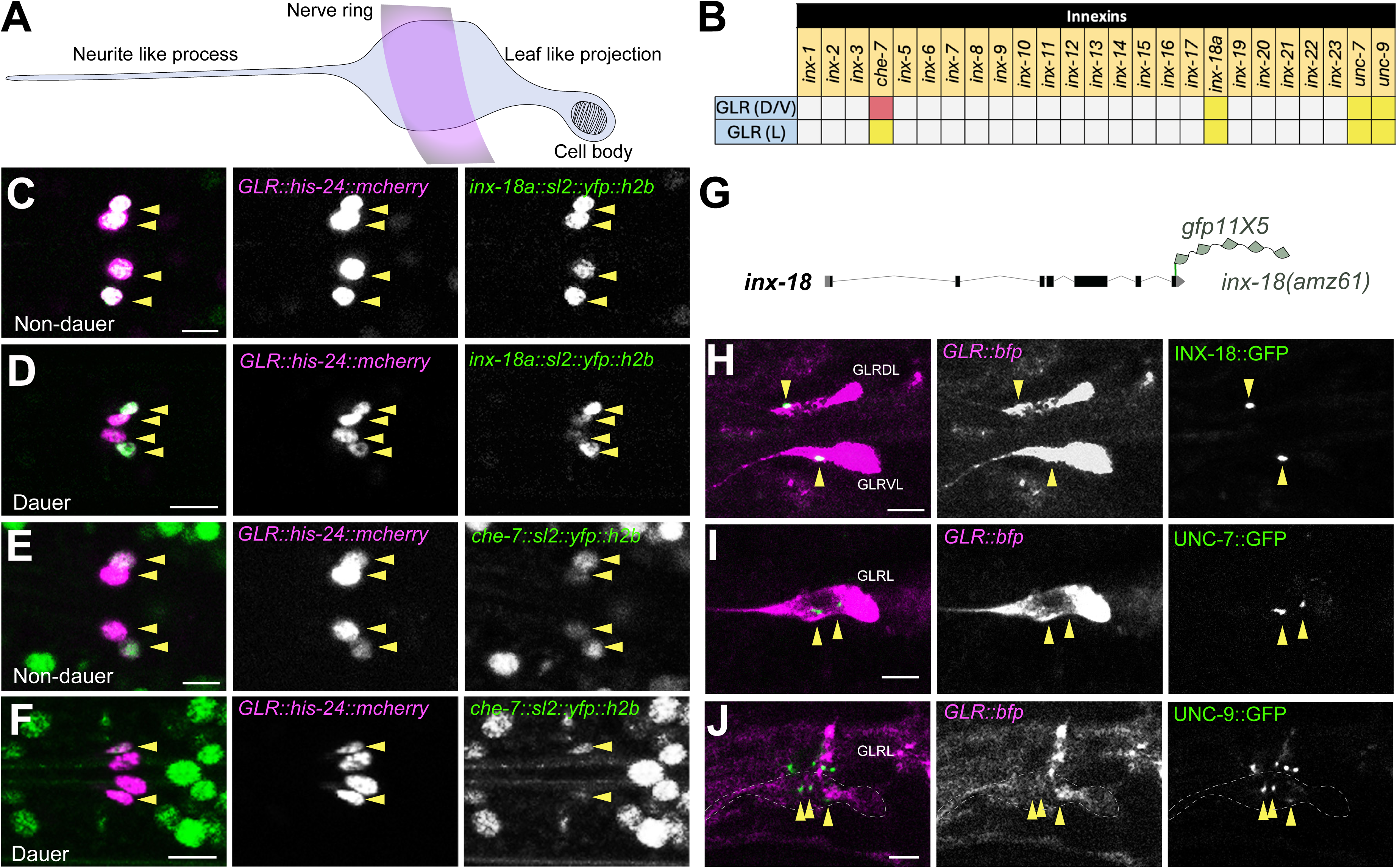
GLR glial cells express four innexins. (A) Schematic representation of a GLR glial cell (in grey). *C. elegans* nerve ring position is marked by a magenta band. (B) Expression of innexin genes in GLR glial cells. Yellow indicates expression of a particular innexin in both non-dauer and dauer-stage animals, whereas red indicates expression only in non-dauer-stage animals. (C-D) The *inx-18* fosmid reporter (*otIs771[inx-18a::SL2::NLS::yfp::H2B)]*) is expressed in all GLR cells in both non-dauer and dauer-stage animals. GLR cell nuclei are marked by the *nep-2p7::his-24::mcherry* reporter strain. (E-F) Expression of the *che-7* reporter allele (*che-7(syb4693[che-7::SL2::GFP::H2B])*) is present in all GLR cells in non-dauers, but is selectively turned off in GLRDL/R and GLRVL/R in dauer animals, while continuing to be expressed in lateral GLRL/R cells. (G) Schematic showing the 5Xsplit-GFP-based *inx-18* endogenous reporter allele, *inx-18a(amz61[inx-18a::5xgfp11])*. (H) Punctate localization of endogenously split-GFP-tagged INX-18 (yellow arrowheads) within the leaf-like projections of dorsal and ventral GLR glial cells, marked by BFP expression (*nep2p7::gfp1-10::t2a::ebfp2*), pseudocoloured in magenta. (I-J) Punctate localization of endogenously split-GFP-tagged UNC-7 and UNC-9 (yellow arrowheads) within the leaf-like projection of lateral GLR glial cells, marked by BFP expression (*nep2p7::gfp1-10::t2a::ebfp2*), pseudocoloured in magenta. Scale bar = 5 μM. Also see Figure S1.

In this study, we demonstrate the molecular heterogeneity of innexin channels within GLR glia and uncover distinct aspects of their function. We identified two distinct channel types formed by innexins INX-18 and UNC-7/UNC-9, which localize to spatially separated compartments and function independently. UNC-7/UNC-9 function as gap junction channels, as has previously been shown to connect GLR cells and RME neurons^9^, whereas INX-18 functions as undocked hemichannels in GLR glia. In this configuration, UNC-7/UNC-9 channels, but not INX-18 hemichannels, regulate Synaptobrevin/SNB-1 localization in RME neurons, as previously reported^9^. However, both channel types function non-redundantly to maintain GLR morphology, potentially by modulating intracellular calcium levels and the activity of the downstream calcium-dependent protease Calpain/CLP-4 and cyclin-dependent kinase CDK-5. In contrast, undocked INX-18 hemichannels, but not UNC-7 channels, cell-autonomously regulate recovery of motility after paralysis induced by exposure to high-salt conditions, potentially in conjunction with tyramine signaling. These findings highlight that glial cells can utilize distinct gap junction and undocked hemichannels to maintain specific aspects of glial morphology and function, thereby regulating animal behaviour.

## RESULTS

### Innexin expression atlas of GLR glial cells

To comprehensively understand the roles of gap junction components in the GLR glia, we characterized the expression of all innexins in these cells, using fosmid-based transcriptional reporter transgenes or CRISPR-based transcriptional reporter alleles^28,29^. A previous study showed that UNC-7 and UNC-9 are expressed in GLR glia, where they form gap junctions between GLR cells and RME motor neurons to regulate RME axon specification^9^. Based on previous studies, *inx-3, inx-6, inx-20, eat-5* (which are expressed in the pharynx and specific neuron-types)^28,30^, *inx-15, inx-16,* and *inx-17* (expressed in the gut)^28^, *inx-21, inx-22* (expressed specifically in the gonad)^31^ were not expressed in the spatial region where GLR cells are present and thus were not included in the analysis. We found that innexins *inx-18* and *che-7,* are expressed in the GLR glial cells during larval and adult stages under replete conditions **(Figure 1B,C,E)**. Our findings are also in concordance with the available RNA sequencing dataset^6^. This combinatorial innexin expression code in GLR cells is distinct from any combination previously observed in neuronal classes and in other glial cells derived from ectodermal lineages^26,28^. Moreover, we found that the GLR-specific combinatorial innexin expression changes when animals enter the developmental diapause stage, called the dauer stage, under adverse environmental conditions^32,33^. We found that during dauer diapause *inx-18* expression remained unaltered, while *che-7* expression was specifically turned off only in dorsal and ventral GLR cells **(Figure 1D,F)**, whereas lateral GLRs retained *che-7* expression. It was shown that the neuronal innexin expression atlas is markedly altered in a neuron-type-specific manner as animals develop into dauer diapause^28^. This suggests that developmental plasticity in innexin expression is not limited to neurons, but also occurs in glial cells.

### Distinct Innexins form Molecularly Independent Channels on GLR leaves

Each of the six GLR glial cells extends flat, leaf-like processes underneath the *C. elegans* nerve ring (the brain neuropil), making contacts with multiple cell types and partitioning the nerve ring from the pharynx **(Figure 1A)**^27,34^. Previous studies have reported functional gap-junction-mediated communication between GLR glia and RME neurons, using UNC-7 and UNC-9^9,34^. To understand how GLR glia utilizes different innexins to form channels, we looked at the localizations of innexins in GLR glia using endogenous fluorescent protein-tagged reporter alleles. To cell-specifically label CHE-7, INX-18, UNC-7 and UNC-9, we generated split-GFP-based^35^ reporter alleles of *che-7, che-7(amz33[che-7::5xgfp11]),* and *inx-18, inx-18(amz61[inx-18::5xgfp11]),* and used previously published alleles of *unc-7* [*unc-7(amz15[unc-7::5xgfp11])*^36^ and *unc-9 [unc-9(miz81[unc-9::splitGFP(11X7)_LoxP])*^37^, where a cassette containing tandem repeats of 11^th^ beta-strand of GFP (*sfGFP11*) were inserted at the C-terminus of these gene loci **(Figure 1G)**. By expressing the GFP1-10 fragment specifically in the GLR glia, we observed INX-18 localization at distinct anterior foci on the leaf-like projections of GLRs **(Figure 1H)**, whereas UNC-7 and UNC-9 localized to discrete puncta on the leaf-like projections **(Figure 1I,J)**. Using a similar cell-specific labeling strategy, we found that the UNC-9 channel-associated stomatin-family regulatory protein, UNC-1, *(amz22[sfGFP(11X5)::unc-1])*, also localizes on the leaf-like projections, similar to UNC-7 and UNC-9 puncta **(Figure S1A,B)**. Using a similar split-fluorophore technique, we did not observe punctate localization of CHE-7 on GLR glial cells, although pan-neuronal expression of the GFP1-10 fragment reconstituted abundant CHE-7 puncta in the nerve ring **(Figure S1A,C,D)**. This suggests that either CHE-7 is not utilized to form channels in GLR glia, or it is localized in a dispersed manner within GLR glia and is undetectable as punctate foci.

To further assess whether these innexins function together or independently to form channels on GLR cells, we tested whether their characteristic localization patterns depend on the presence of co-expressed innexins. Our results suggested that localization of INX-18 remained unaffected in *unc-7(e5)* or *unc-9(e101)* mutant animals **(Figure 2A-D)**. Similarly, we found that UNC-7 localization remained unaffected in *inx-18(ok2454)* mutant animals **(Figure 2E,F)**, suggesting that these innexins assemble and localize independently.

**Figure 2:**
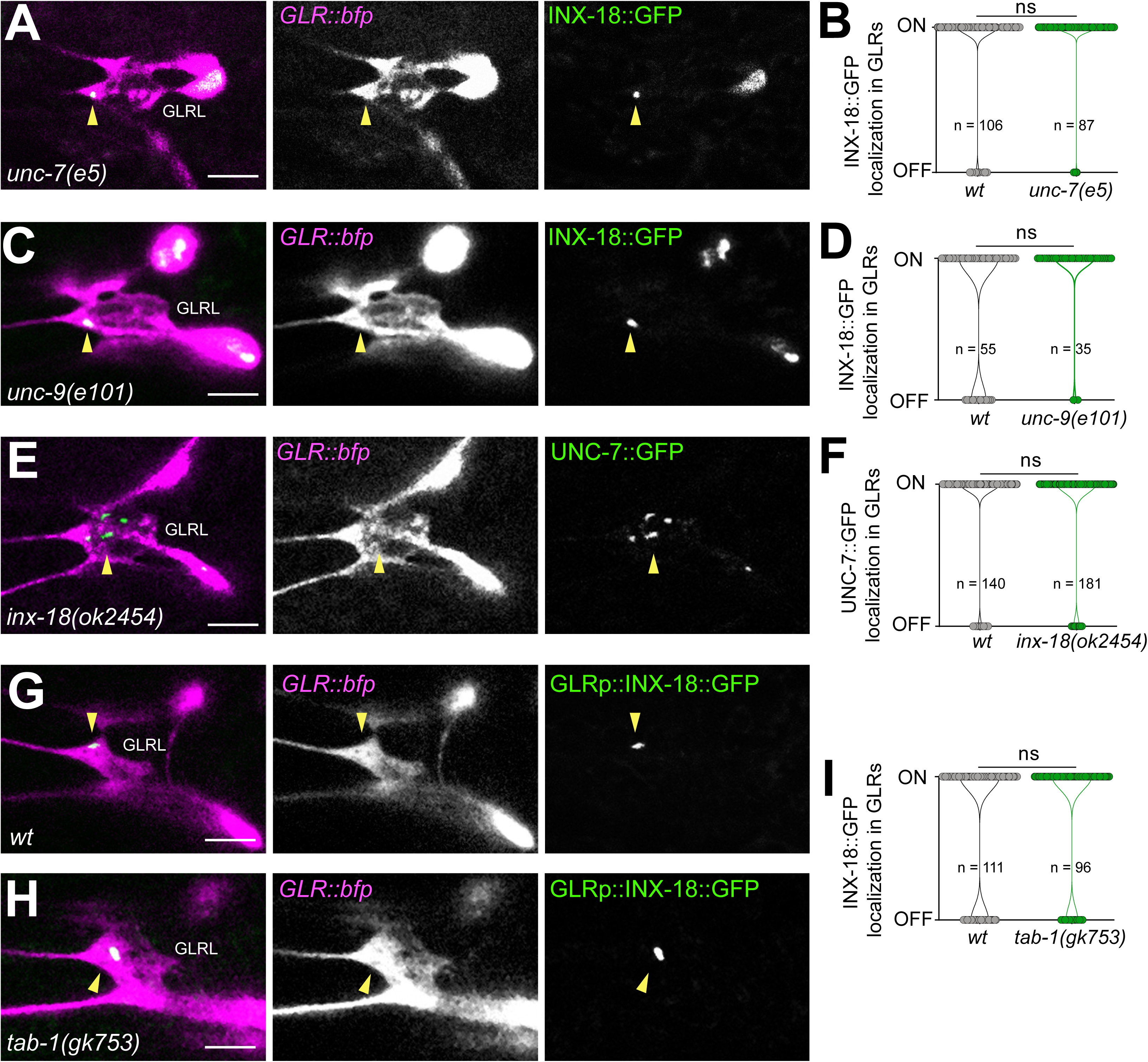
INX-18 and UNC-7 localizations within GLR cells are independent of each other. GLR cells are marked by BFP expression, pseudocoloured in magenta. Scale bar = 5 mM. (A and C) Punctate localization of endogenously split-GFP-tagged INX-18 (yellow arrowheads) in*, inx-18a(amz61[inx-18a::5xgfp11])* within GLR cells (pseudocoloured in magenta) remained intact in *unc-7(e5)* and *unc-9(e101)* mutant animals, respectively. (B and D) Violin plots showing quantifications of the data shown in Panels A and C, respectively. Statistical comparisons between groups were performed using Kolmogorov-Smirnov test. ns=non-significant. n=number of glial cells, specified beneath each distribution. (E) Punctate localization of endogenously split-GFP-tagged UNC-7 (yellow arrowheads) in *unc-7(amz15[unc-7::5xgfp11])*, within GLR cells (pseudocoloured in magenta), remained intact in *inx-18(ok2454)* mutant animals. (F) Violin plots show quantifications of the data shown in Panel E. Statistical comparisons between groups were performed using Kolmogorov-Smirnov test. ns=non-significant. n=number of glial cells, specified beneath each distribution. (G) GLR-specific expression of GFP-tagged INX-18 in *nep-2p7::inx-18a::gfp* shows punctate localization (yellow arrowheads) within the leaf-like projection in lateral glial cells (pseudocoloured in magenta). (H) Punctate localization of GFP-tagged INX-18 (yellow arrowheads) in *nep-2p7::inx-18a::gfp* within the leaf-like projection in lateral GLR glial cells (pseudocoloured in magenta) was not disrupted in *tab-1(gk753)* mutant animals. (I) Violin plots show quantifications of the data shown in Panels G and H. Statistical comparisons between groups were performed using Kolmogorov-Smirnov test. ns=non-significant. n=number of glial cells, specified beneath each distribution.

The leaf-like projections of the lateral GLR cells extensively overlap with the lateral RMEL/R neurons and form gap junctions, as inferred from the serial-section electron micrograph (EM) reconstruction^27,34^. Previously, it was shown that UNC-7 and UNC-9, along with the associated Stomatin-family member, UNC-1, form gap junctions between GLR glia and RME neurons^9^. To determine whether INX-18 also forms gap junctions between GLR and RME, we examined INX-18 puncta on lateral GLR cells in animals mutant for the homeodomain transcription factor *tab-1(gk753),* an ortholog of the vertebrate Bsx protein^38^, in which the partnering lateral RME neurons fail to differentiate^39^. We found that the punctate localization of INX-18 on lateral GLR cells remained unaltered in the absence of partnering lateral RME neurons **(Figure 2G-I)**. These results suggest that GLR cells partition distinct innexins into discrete zones along their leaf-like projections and may utilize them to regulate different aspects of GLR function.

### Distinct innexins regulate specific aspects of GLR glia function and morphology

To understand the functional significance of innexin channels in GLR glia, we examined how innexin mutations affect GLR glial function. GLR cells have been shown to regulate sensitivity to high salt concentrations, putatively by maintaining osmotic homeostasis with the external surroundings^6^. Wild-type animals, when exposed to high concentrations of sodium chloride (NaCl), become paralyzed within a short period but recover quickly^6^. On the other hand, GLR-ablated animals remain paralysed for extended periods of time under identical conditions^6^. We found that *inx-18(ok2454)* and *unc-7(e5)* mutant animals also exhibit prolonged paralysis compared to wild-type animals upon exposure to 200 mM NaCl, even though the GLR cells are intact on their own **(Figure 3A)**. We did not find any significant effect on salt hypersensitivity behaviour in *che-7(ok2373)* mutant animals **(Figure S2A).** However, both *inx-18* and *unc-7* are expressed in other nervous system cell types. To understand whether innexin channels function cell-autonomously to regulate the salt-hypersensitivity behaviour, we deleted *inx-18* and *unc-7* specifically in GLR cells using a loxP-recombination-site-flanked conditional knockout allele of *inx-18*(*amz71amz78 [loxP::inx-18::loxP]*) **(Figure S2B)** and *unc-7(amz14ot895[loxP::unc-7::tagRFP-t::loxP])*^36^, respectively. GLR cell-specific loss of *inx-18,* by expression of Cre recombinase specifically in these cells, affected the salt-hypersensitivity response similar to systemic *inx-18(ok2454)* mutants **(Figure 3B, S2C)**. However, unlike the *unc-7(e5)* mutation, GLR cell-specific loss of *unc-7* had no significant effect on the salt-hypersensitivity response of the animal **(Figure S2D)**. These results suggest that INX-18 channels specifically in GLR cells regulate the salt-hypersensitivity behaviour, while UNC-7 functions non-cell autonomously.

**Figure 3:**
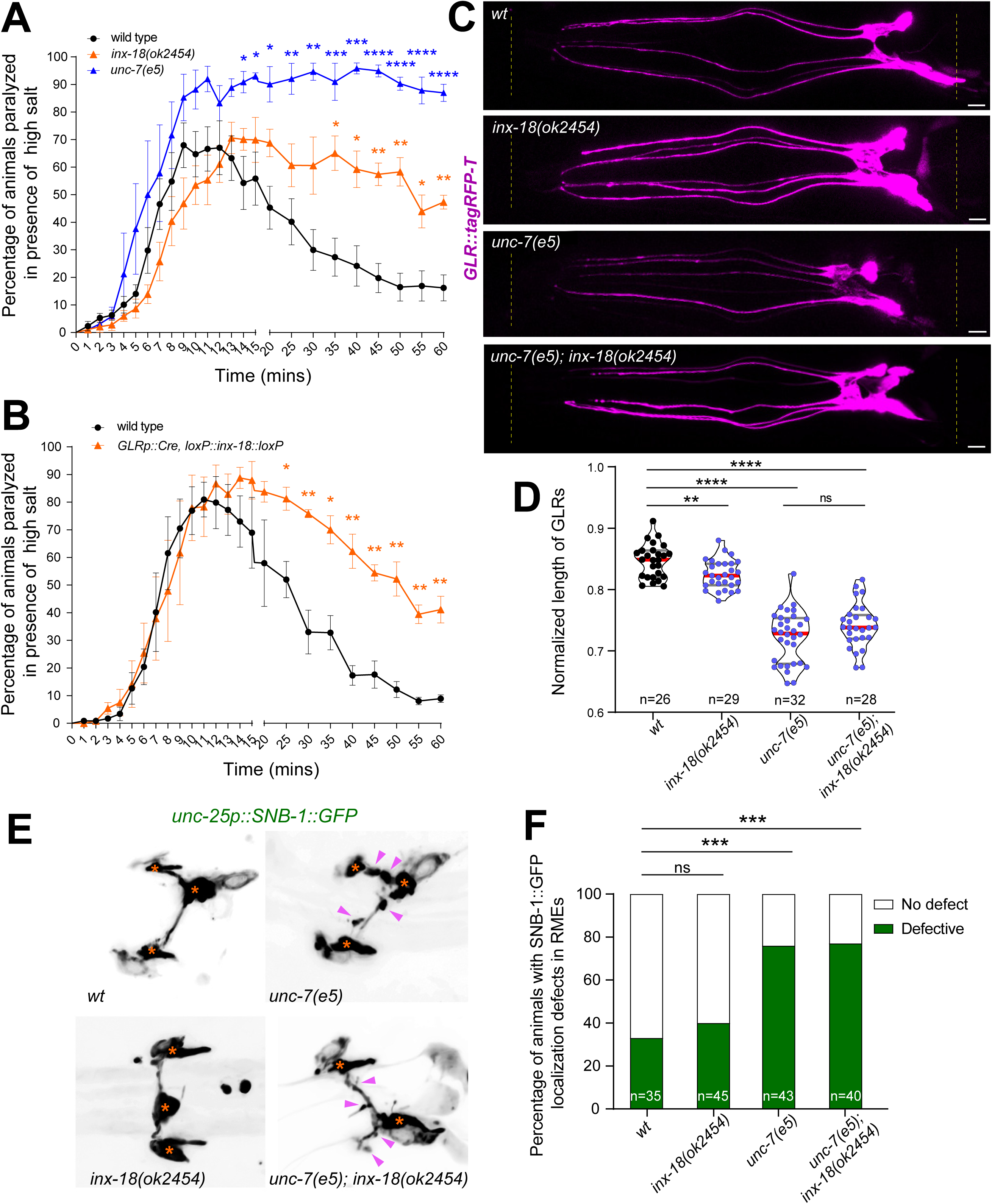
INX-18 and UNC-7 regulate GLR glia morphology and functions. (A and B) Statistical comparisons between groups were performed using multiple t-tests followed by two-stage step-up procedure of Benjamini, Krieger and Yekutieli correction, with FDR(Q)=5%. Distributions shown are standard error of mean. *p<0.05, **p<0.01, ***p<0.001, ****p<0.0001 – compared to wild-type animals. (A) *inx-18(ok2454)* and *unc-7(e5)* mutant animals undergo paralysis when exposed to 200 mM NaCl, similar to wild-type animals. However, these mutants show significantly delayed recovery of motility than wild-type animals. (B) Cre recombinase-mediated, GLR-specific deletion of *inx-18* in loxP-flanked *inx-18* animals, *inx-18a(amz71amz78[loxP::inx-18a::loxP])*, significantly affected post-paralysis recovery of motility when exposed to 200 mM NaCl. (C) Animals mutant for either *inx-18(ok2454)* or *unc-7(e5)*, or double mutants for *unc-7(e5); inx-18(ok2454)*, had significantly shorter GLR glia length (pseudocoloured in magenta). GLR lengths were normalized to the distance between the nose tip of the animal and the grinder in the posterior pharynx (indicated by the yellow dotted vertical lines) within the same animal. Scale bar=5 μm. (D) Violin plots showing quantifications of the data shown in Panel C. Each point in the violin plot indicates the normalized average length of GLRs within a single animal. Red horizontal line denotes the median, and grey lines indicate quartiles. Statistical comparisons between groups were performed using unpaired t-test with Welch’s correction. n=number of animals, specified beneath each distribution. ns=non-significant, *p<0.05, **p<0.01, ***p<0.001, ****p<0.0001. (E) Representative images showing expression of SNB-1::GFP within RME neurons. Cell bodies are marked by orange asterisks. Animals mutant for either *unc-7(e5)* or double mutants for *unc-7(e5); inx-18(ok2454)* show significantly higher abnormal punctate accumulation of SNB-1::GFP within RME neurites (marked by magenta arrowheads). (F) Stacked bar plots representing data shown in Panel E. Each bar represents the percentage of animals showing SNB-1::GFP accumulation in RME in each genotype. Statistical comparisons between indicated groups were performed using Fisher’s exact test; ns=non-significant, ***p<0.001. n=number of animals. Also see Figure S2.

Innexins and connexins have been shown to be associated with the maintenance of glia morphology^10,40^. We found that *inx-18* and *unc-7* regulate the length of GLR glia cells (**Figure 3C,D)**. We found only a mild effect on GLR morphology in *che-7(ok2373)* mutant animals **(Figure S2E).** The effect on GLR morphology was most severe in *unc-7(e5),* however, we did not observe an enhanced defect in *unc-7(e5); inx-18(ok2454)* double mutant animals, suggesting a non-redundant role for *unc-7* and *inx-18* in regulating GLR glia morphology **(Figure 3C,D)**. GLR cells are anatomically positioned beneath the nerve ring, the brain neuropil of the animal, and have been proposed to function as supporting buttresses. Ablation of GLR glia precursors has been shown to cause anterior displacement of the nerve ring^41^. To understand whether shortening of the GLR glia had any associated defects in the nerve ring positioning, we examined the relative positioning of RME motor neurons, which are suggested to form the peripheral boundary of the developing nerve ring in the embryo^27^. We observed a significant anterior shift in the nerve ring position in *unc-7(e5)* mutants, but not in *inx-18(ok2454)* mutants **(Figure S2F)**, further suggesting distinct roles for UNC-7 and INX-18 in regulating GLR glia-associated function.

In addition to regulating GLR morphology, gap junctions formed by UNC-7 and UNC-9 between GLR glia and RME neurons, along with associated UNC-1, affect RME axon specification, resulting in aberrant localization of the presynaptic protein Synaptobrevin (SNB-1) within RME axons^9^ **(Figure 3E,F)**. We found that INX-18, which does not interact with RME neurons, also does not affect SNB-1 localization within RME axons **(Figure 3E,F)**, suggesting that GLR glial cells utilize molecularly distinct innexin channels to perform specific, non-overlapping functions.

### INX-18 functions as undocked hemichannels to regulate GLR glia functions along with UNC-7 gap junctions

To understand the mechanism by which INX-18 regulates GLR functions, we first tried to understand the nature of INX-18 channels in GLR cells. We found that INX-18::GFP localization in GLR cells remained unaltered in *inx-18(ok2454)* mutant animals **(Figure S3A,B),** suggesting INX-18 does not form homotypic gap junction channels in GLRs. We also found that INX-18 localization on GLR cells is independent of other GLR-expressed innexins *unc-7* and *unc-9* **(Figure 2A,B)**. INX-18 localization in GLRs also remained unaffected in the absence of *inx-1* and *inx-7*, innexins that are expressed in neurons which form gap junctions with GLR glial cells according to the EM reconstruction studies^27,28,34,42^ **(Figure S3C-F)**.

Innexins and their homologous pannexin proteins in vertebrates have also been shown to function as undocked hemichannels across cell types^19,20,43,44^. To determine whether INX-18 functions as undocked hemichannels in GLR glial cells, independent of any docking innexin partner, we generated a chimeric INX-18 protein that would lose the ability to form gap junction channels but retain the ability to form and function as undocked hemichannels **(Figure 4A, S3G)**. It has been shown that replacing the extracellular loop 2 (ECL2) of UNC-7 with that of mouse Pannexin1 (mPANX1) arrested chimeric UNC-7 channels as undocked hemichannels while retaining channel properties^44^. We found that GLR-specific expression of the chimeric INX-18-PANX1-ECL2::GFP exhibited punctate localization at a location characteristic of endogenous INX-18 puncta on the GLR leaf-like projection in *inx-18(ok2454)* mutant animals **(Figure 4B)**. Moreover, we found that GLR-specific expression of the chimeric INX-18-PANX1-ECL2 was sufficient to rescue both the salt-hypersensitivity and GLR morphology defect associated with the *inx-18(ok2454)* mutant animals **(Figure 4C-E)**, suggesting that INX-18 functions as undocked hemichannels in GLR glial cells.

**Figure 4:**
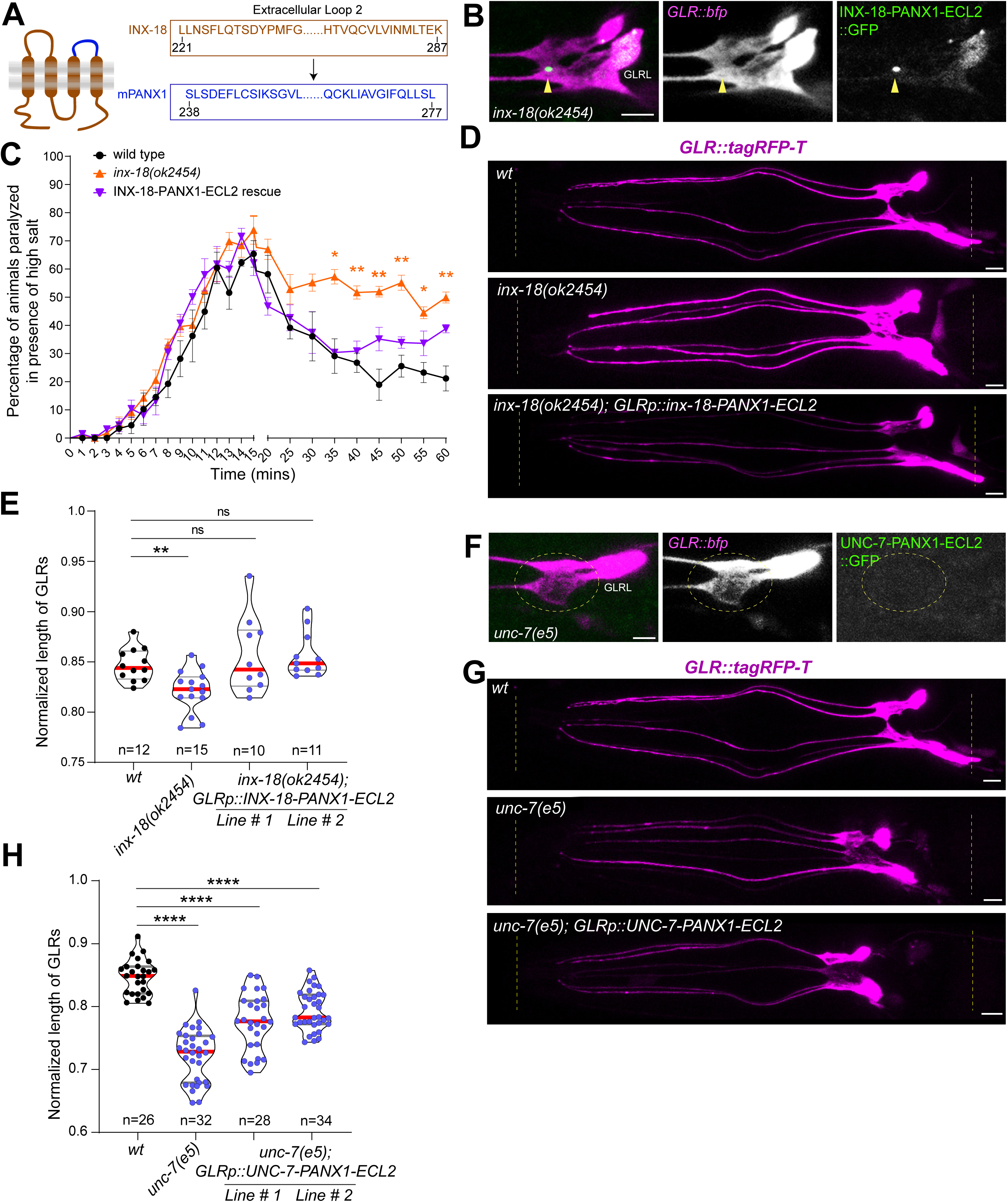
INX-18 functions as undocked hemichannels, while UNC-7 functions as gap junctions. (A) Left: Schematic showing the structure of the chimeric INX-18-PANX1-ECL2, in which the extracellular loop 2 (ECL2) of INX-18 (in brown) was replaced with that of mouse Pannexin1 (mPANX1) (shown in blue). Right: Sequence of INX-18 ECL2 (in brown) and mPANX1 ECL2 (in blue). Numbers indicate the corresponding amino acid positions. (B) GLR-specific expression of GFP-tagged INX-18-PANX1-ECL2 in *nep-2p7::inx-18-mPANX1-ECL2::gfp* results in punctate localization (yellow arrowheads) within the leaf-like projection in lateral glial cells (pseudocoloured in magenta), similar to that of endogenous INX-18 (Figure 1H, 2G). (C) GLR-specific expression of chimeric INX-18-PANX1-ECL2 in *inx-18(ok2454)* mutant animals (Transgenic line # 2) was sufficient to rescue the delayed recovery of motility following paralysis when exposed to 200 mM NaCl, reminiscent of wild-type animals. Statistical comparisons between groups at each time point were performed using multiple t-tests followed by two-stage step-up procedure of Benjamini, Krieger and Yekutieli correction, with FDR(Q)=5%. Distributions shown are standard error of mean. *p<0.05, **p<0.01, ***p<0.001, ****p<0.0001 – compared to wild-type animals. (D) Representative images showing that GLR-specific expression of chimeric INX-18-PANX1-ECL2 in *inx-18(ok2454)* mutant animals was sufficient to rescue the GLR glia (pseudocoloured in magenta) length observed in the *inx-18(ok2454)* animals. GLR lengths were normalized to the distance between the nose tip of the animal and the grinder in the posterior pharynx (indicated by the yellow dotted vertical lines) within the same animal. Representative images for *wt* and *inx-18(ok2454)* have been reused from Figure 3C. Scale bar=5 μm. (E) Violin plots showing quantifications of the data shown in Panel D. Each point in the violin plot indicates the normalized average length of GLRs within a single animal. Red horizontal line denotes the median, and grey lines indicate quartiles. Statistical comparisons between groups were performed using unpaired t-test with Welch’s correction for all, except for Kolmogorov-Smirnov test used for Line#2 of GLRp::INX-18-PANX1-ECL2 following a non-parametric distribution. n=number of animals, specified beneath each distribution. ns=non-significant, *p<0.05, **p<0.01, ***p<0.001, ****p<0.0001. (F) GLR-specific expression of GFP-tagged UNC-7-PANX1-ECL2 in *nep-2p7::unc-7-mPANX1-ECL2::gfp* did not show punctate localization (yellow dotted circle) within the leaf-like projection in lateral glial cells (pseudocoloured in magenta), unlike the localization of endogenous UNC-7 (Figure 1I). (G) Representative images showing that GLR-specific expression of the chimeric UNC-7-PANX1-ECL2 in *unc-7(e5)* mutant animals does not rescue the GLR glia (pseudocoloured in magenta) length observed in the *unc-7(e5)* animals. GLR lengths were normalized to the distance between the nose tip of the animal and the grinder in the posterior pharynx (indicated by the yellow dotted vertical lines) within the same animal. Representative images for *wt* and *unc-7(e5)* have been reused from Figure 3C. Scale bar=5 μm. (H) Violin plots show quantifications of the data shown in Panel G. Each point in the violin plot indicates the normalized average length of GLRs within a single animal. Red horizontal line denotes the median, and grey lines indicate quartiles. Statistical comparisons between different groups were performed using unpaired t-test with Welch’s correction. n=number of animals, specified beneath each distribution. ns=non-significant, *p<0.05, **p<0.01, ***p<0.001, ****p<0.0001. Also see Figure S3.

UNC-7 was shown to function as undocked hemichannels in thermosensory AFD neurons to regulate thermotaxis behavior^44^. We tested whether UNC-7 also functions as undocked hemichannels in GLRs to regulate GLR morphology, in addition to its known role in forming gap junction channels between GLR glia and RME neurons. GLR cell-specific expression of the chimeric UNC-7-PANX1-ECL2 in *unc-7(e5)* mutant animals only partially rescued GLR glial cell morphology defects **(Figure 4G,H)**. Similarly, GLR cell-specific expression of the chimeric UNC-7-PANX1-ECL2 failed to rescue the salt-hypersensitivity defects observed in *unc-7(e5)* mutant animals **(Figure S3H),** further supporting the non-cell-autonomous role of UNC-7 in regulating the salt-hypersensitivity behaviour. Moreover, unlike chimeric INX-18-PANX1-ECL2::GFP, chimeric UNC-7-PANX1-ECL2::GFP did not exhibit punctate localization in the leaf-like projections of GLR cells **(Figure 4F)**, potentially due to fewer UNC-7 hemichannels rendering it undetectable as distinct puncta. These results suggest that different innexins expressed in the same glial cell can assemble in distinct mechanisms to regulate cell functions.

### Innexin channels regulate GLR glia morphology downstream of calcium signalling in GLRs

Undocked hemichannels and gap junctions across cell types and species are intricately linked to intracellular calcium ion levels^9,16,45^. Previously, it was shown that GLR-RME gap junctions regulate RME axon specification by modulating intracellular calcium levels in RME neurons^9^. To determine how undocked INX-18 hemichannels and UNC-7 gap junction channels in GLR cells regulate glial functions, we downregulated intracellular calcium levels using a cell-permeable calcium chelator, BAPTA-AM. We found that reducing accessible calcium ions with BAPTA-AM significantly decreased the length of GLR cells in otherwise wild-type animals **(Figure 5A,B)**. Furthermore, treatment of *inx-18(ok2454)* animals with BAPTA-AM did not further enhance the GLR-length phenotype **(Figure 5C,D)**, suggesting INX-18 hemichannels and intracellular calcium levels regulate GLR morphology in a non-redundant manner. BAPTA-AM treatment reduces intracellular calcium at the systemic level. To determine whether intracellular free calcium levels in GLR glia cell-autonomously regulate cellular morphology, we expressed a version of the vertebrate calcium-buffer protein calbindin D28K^46^ specifically in GLR cells. Expression of calbindin D28K in *C. elegans* has been shown to sequester free calcium in a cell-specific manner, thereby lowering its levels in specific cell types. We found that, similar to BAPTA-AM treatment, GLR-specific expression of calbindin D28K significantly reduced GLR glia length in otherwise wild-type animals, reminiscent of innexin mutant animals **(Figure 5E,F)**.

**Figure 5:**
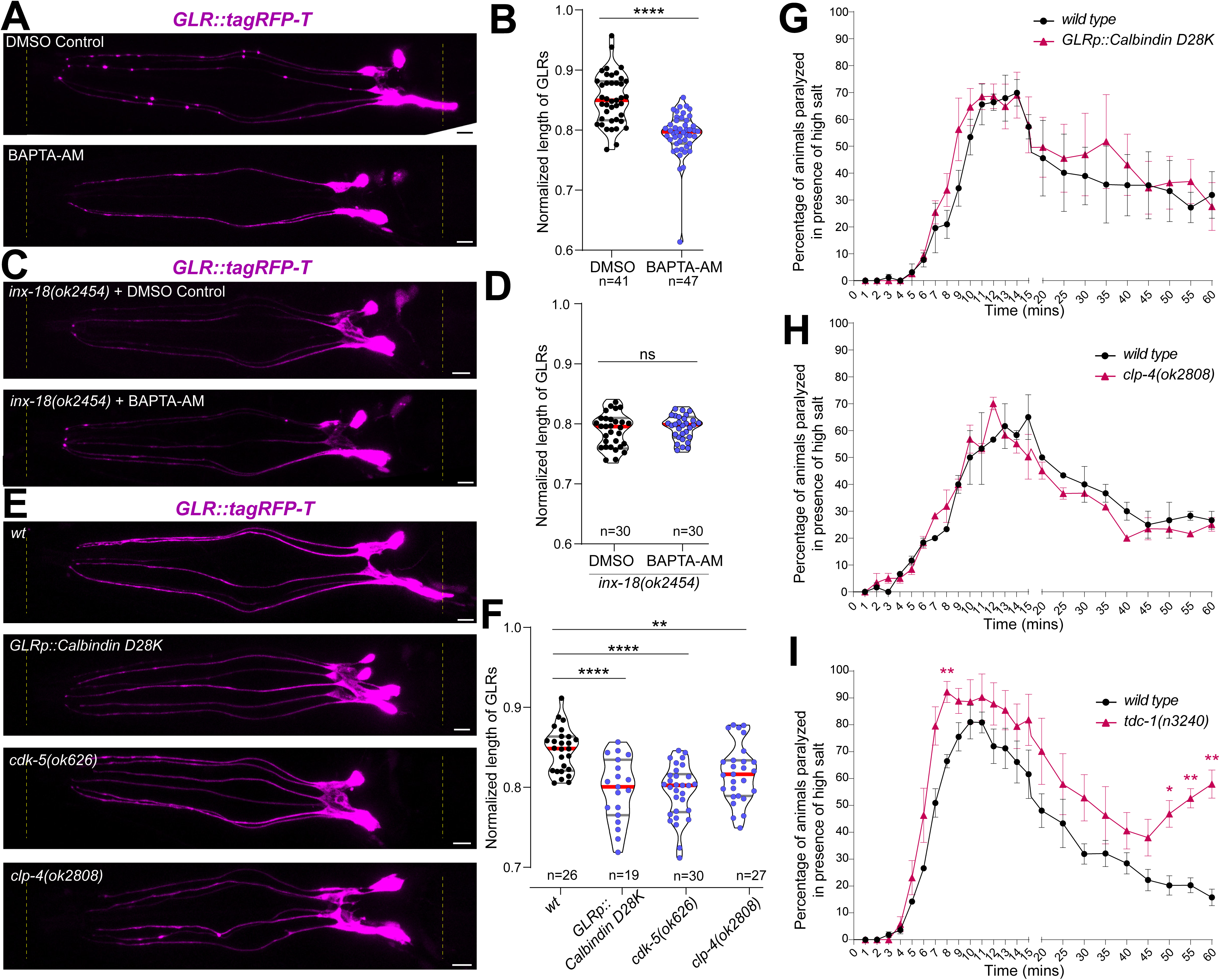
Ca^2+^-signalling regulates GLR glia morphology, but not salt hypersensitivity behaviour. (A,C,E) GLR glia projections, pseudocoloured in magenta. The distance between the nose tip of the animal and the grinder in the posterior pharynx (indicated by yellow dotted vertical lines) within the same animal. Scale bar=5 μm. (B,D,F) Each point in the violin plot represents the normalized average length of GLRs within a single animal. Red horizontal line denote the median, and grey lines indicate the quartiles. n=number of animals, specified beneath each distribution. (A) Representative images showing that treatment with BAPTA-AM significantly shortens the length of GLR glia compared to control, DMSO-treated animals. (B) Violin plots show quantifications of the data shown in Panel A. Statistical comparisons between groups were performed using Kolmogorov-Smirnov test. ****p<0.0001. (C) Representative images showing that treatment with BAPTA-AM does not further affect the length of GLR glia in *inx-18(ok2454)* mutant animals, compared to control, DMSO-treated animals of the same genotype. (D) Violin plots show quantifications of the data shown in Panel C. Statistical comparisons between groups were performed using unpaired t-test with Welch’s correction. ns=non-significant. (E) Representative images showing that GLR-specific expression of human Calbindin D28K (*nep-2p7::Calbindin D28K*) and mutations in either *cdk-5(ok626)* or *clp-4(ok2808)* significantly shortens GLR glia length. Representative image for *wt* has been reused from Figure 3C. (F) Violin plots show quantifications of the data shown in Panel E. Statistical comparisons between groups were performed using unpaired t-test with Welch’s correction. **p<0.01, ****p<0.0001. (G,H,I) Statistical comparisons were performed using multiple t-tests followed by two-stage step-up procedure of Benjamini, Krieger and Yekutieli correction, with FDR(Q)=5%. Distributions shown are standard error of mean. *p<0.05, **p<0.01 – compared to wild-type animals. (G-H) GLR-specific expression of human Calbindin D28K (*nep-2p7::Calbindin D28K*) and mutations in *clp-4(ok2808)* do not affect paralysis or post-paralysis recovery of motility when exposed to 200 mM NaCl, compared with wild-type animals. (I) *tdc-1(n3240)* mutant animals undergo paralysis when exposed to 225 mM NaCl, earlier than wild-type animals. However, contrary to wild-type animals, *tdc-1* mutants show failure in paralysis recovery after 60 minutes. Also see Figure S4.

In RME neurons, regulation of intracellular calcium concentration by GLR-RME gap junctions activates the calcium-dependent cysteine protease Calpain/CLP-4 and the downstream cyclin-dependent kinase CDK-5, which in turn affects microtubule organization and axon specification in RME neurons^9^. To understand how intracellular calcium levels regulated by innexin channels control GLR glia morphology, we tested the involvement of *clp-4* and *cdk-5.* We found that both *clp-4(ok2808)* and *cdk-5(ok626)* mutant animals exhibited significantly reduced GLR glia length, reminiscent of the phenotype observed in animals expressing calbindin D28K in GLR cells and in innexin mutants **(Figure 5E,F**; **3D).** These results suggest that innexin channels may regulate intracellular calcium levels in GLR glial cells, which, in a CLP-4- and CDK-5-dependent manner, regulate glial morphology.

### Salt-hypersensitivity behaviour is regulated by a calcium level-independent mechanism

To understand whether innexin channels regulate salt-hypersensitivity behaviour also through the same mechanism, we tested animals that were either mutant for *clp-4(ok2808)* and *cdk-5(ok626),* or transgenic animals overexpressing calbindin D28K specifically in GLR cells. Animals treated with the calcium chelator BAPTA-AM showed reduced movement and were not assayed for high-salt-induced paralysis. Interestingly, we found that all *GLRp::calbindin D28K-expressing, cdk-5(ok626),* and *clp-4(ok2808)* mutant animals showed unaltered salt-hypersensitivity behavior, comparable to wild-type animals **(Figure 5G,H, S4A)**. These results suggest that innexin channels regulate GLR glia-mediated salt-hypersensitivity behaviour independently of intracellular calcium levels and regulate glia morphology through distinct, calcium level-independent molecular mechanisms.

Undocked hemichannels and gap junctions are also linked to the regulation of cyclic nucleotide (cAMP and cGMP) levels in the cell^47–49^. To test whether INX-18 undocked hemichannels regulate cAMP and/or cGMP levels to modulate salt-hypersensitivity behaviour, we examined animals mutant for cyclic nucleotide phosphodiesterases (PDEs), which break down cAMP and/or cGMP and are major determinants of cellular cyclic nucleotide levels. The *C. elegans* genome contains six PDEs (*pde-1* through *pde-6*). Out of these, - *pde-4* expression is significantly enriched in GLR cells^6^. We examined GLR glia morphology and salt-hypersensitivity behaviour in *pde-4(ce268)* mutant animals, which may alter cyclic nucleotide levels. Our data suggested that GLR morphology and salt-hypersensitivity behaviour remained unaffected in the absence of *pde-4* **(Figure S4B,C)**, indicating that innexin channels may function independently of cyclic nucleotide levels.

Pannexin undocked hemichannels have been shown to functionally interact with ligand-gated ion channels, including NMDA receptors^16,23,24^. One of the biogenic amines, tyramine, gates the chloride channel LGC-55, which is particularly enriched in GLR cells^6,50^. Moreover, tyramine functions as a neuromodulator across species, including in *C. elegans*, where it regulates locomotion and head muscle activity^51,52^. To test whether tyramine signaling regulates salt-hypersensitivity behavior, we examined *tdc-1(n3240)* mutant animals, which encode the sole ortholog of the tyrosine decarboxylase enzyme in *C. elegans* and are an essential component of the tyramine biosynthesis pathway^51^. We found that *tdc-1(n3240)* mutant animals exhibited slightly accelerated paralysis under high-salt conditions and continued to show significantly higher paralysis even after 60 minutes or at the end of the assay period. **(Figure 5I)**. Our results suggest that undocked INX-18 hemichannels and UNC-7 gap junction channels in GLR glia may function together with tyramine signaling to regulate salt-hypersensitivity behavior in the animal.

**Figure 6:**
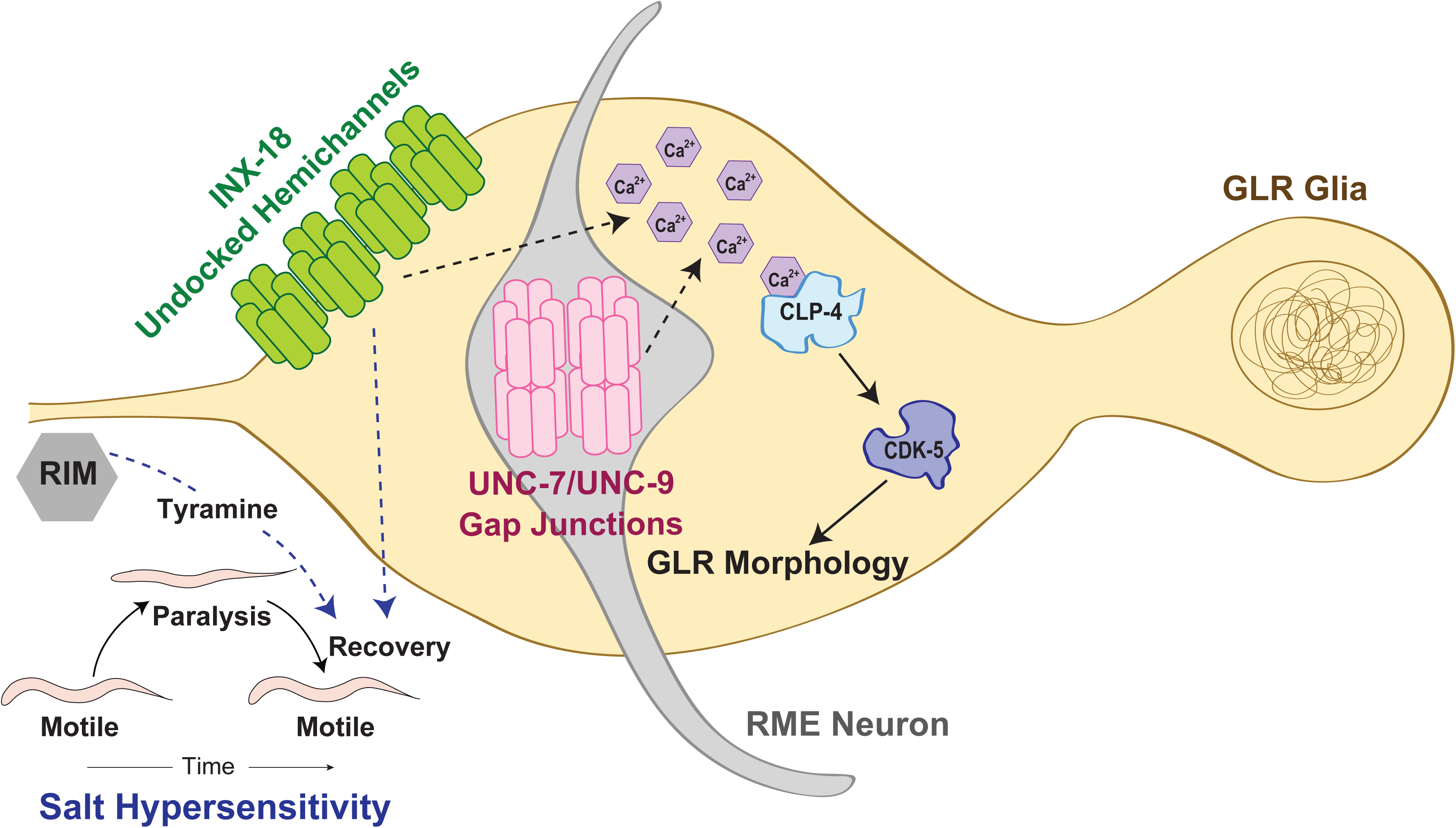
Model showing the distinct molecular organization of innexin channels and their roles in regulating GLR glia function. Model showing the distinct molecular configuration of innexin channels within GLR glial cells. INX-18 forms independent undocked hemichannels, whereas UNC-7 and UNC-9 function predominantly as gap junction channels between GLR and RME neurons. In this configuration, both channel types regulate GLR glial morphology, potentially by modulating intracellular calcium levels and the downstream calcium-dependent protease Calpain/CLP-4 and the cyclin-dependent kinase CDK-5. However, UNC-7-UNC-9 channels specifically regulate Synaptobrevin/SNB-1 localization in RME neurites (synaptic partner of GLRs)^9^, whereas INX-18 hemichannels regulate the salt-hypersensitivity phenotype of the animal, potentially interacting with tyramine signaling.

## DISCUSSION

Glial cells play integral roles in nervous system organization and function, and there is an increasing appreciation for the significance of gap junctions and undocked hemichannels in these cells. Recent transcriptomic studies in both vertebrates and invertebrates have identified expression of multiple innexins and connexins within individual glial cell classes^6,25,26^. However, the utilization of these channel components at the single-cell level remains unclear. Recent evidences suggest that individual neurons in the nervous systems of *C. elegans* and zebrafish, which express multiple innexins and connexins, employ a subset of these proteins to form molecularly distinct gap junction channels that cluster together to form heterochannel electrical synapses^36,53^. In this study, we show that individual GLR glial cells, which share gene expression profiles with vertebrate astrocytes and endothelial cells^6^, express multiple neuronal innexin genes. However, in contrast to neurons, which form heterochannel synapses, GLR glial cells segregate these innexins into distinct domains within their leaf-like processes that separate the nerve ring from the pseudocoelom Furthermore, while UNC-7 and UNC-9 are utilized to form gap junction channels, potentially between GLR cells and RME neurons as previously suggested^9^, INX-18 forms independent undocked hemichannels. Another innexin, CHE-7, does not localize to any foci in GLR cells, despite being heavily transcribed.

Our work showed that INX-18 undocked hemichannels and UNC-7-UNC-9 gap junction channels are localized to specific domains within the leaf-like projections in GLR glial cells. Localization of endogenous innexin hemichannels has not previously been reported. Localization of specific innexin and connexin gap junction channels depends on channel-specific synaptic transport mechanisms, including regulation by particular kinesin motors and cellular cAMP levels^36,54^. Similarly, the turnover of specific channel types relies on channel-specific mechanisms involving cytoskeleton-interacting proteins and atypical kinesin motors^36,55^. Changes in these regulatory mechanisms under altered environmental conditions have also been associated with the plastic changes in innexin localization. Further work is required to elucidate the molecular mechanisms that regulate the distinct localization of INX-18 hemichannels.

UNC-7 is known to function as undocked hemichannels in mechanosensory and thermosensory neurons of *C. elegans*^43,44^. In contrast, INX-18 forms gap junctions within the nociceptive sensory circuit^49^, indicating that these innexins can assemble into both types of channel architectures. Additionally, INX-18 has been shown to form gap junction channels with UNC-9^56^, yet it preferentially forms undocked hemichannels within GLR cells despite the availability of a partner innexin. Context-dependent glycosylation of pannexins to their extracellular loops regulates the formation of undocked hemichannels^57,58^. Despite growing evidence for innexin and connexin hemichannels in vivo, the molecular mechanisms behind their hemichannel formation remain unclear. Our findings establish a foundation for further studies to understand how individual glial cells or neurons decide whether to form gap junctions or hemichannels, and select specific innexin proteins for each purpose. Further work is required to elucidate the functional diversity and biophysical properties of innexin hemichannels.

Our results show that glial gap junctions and undocked hemichannels regulate specific functional and morphological features through distinct cellular mechanisms. Gap junctions and hemichannels regulate GLR glia morphology by modulating calcium signaling through calcium-dependent Calpain/CLP-4 and downstream CDK-5, whereas INX-18 hemichannels specifically regulate high-salt-induced paralysis through distinct mechanisms. Further work is needed to understand whether innexin channels directly mediate Ca^2+^ ion passage or modulate other ion channels to regulate Ca^2+^ ion levels in GLR cells. Furthermore, our findings reveal a functional similarity between innexin channels in GLR cells and tyramine signaling in the regulation of high-salt-induced paralysis behaviour. However, the molecular mechanism underlying this functional interaction remains to be elucidated. Altogether, our work provides a framework for understanding how individual glial cells may utilize the repertoire of channel components to modulate distinct cellular pathways and regulate diverse cellular functions.

## MATERIALS AND METHODS

### *C. elegans* Maintenance and Strain Generation

*C. elegans* were cultured in 60 mm Nematode Growth Media (NGM) petri plates seeded with *E. coli* (OP50) bacteria as a food source at 20-22°C, or 25°C as previously described (Brenner 1974; Stiernagle 2006). *C. elegans* strain N2-Bristol was used as wildtype control (unless otherwise mentioned). To induce dauer arrest, animals were maintained at 25°C under standard conditions of overcrowding, starvation, and higher temperatures, as previously described^32,33^. Dauer were selected by treating them with 1% SDS.

Transgenic strains were generated using the standard microinjection techniques (Evans 2006). Details of all injections and strains used are mentioned in the strain list **(Table S1)**. For genomic integration, the extrachromosomal strain was treated with UV irradiation (Evans 2006).

### CRISPR/Cas9-mediated genome editing

CRISPR/Cas-9 mediated genome editing was done using purified Cas9 (IDT, catalog # 1081059), tracrRNA (IDT, catalog # 1072533), and crRNA (IDT), as per methods described previously (Ghanta and Mello 2020; Eroglu et al. 2023; Vats et al. 2025). Details of all crRNAs and repair oligos or double-stranded DNA fragments are listed below:

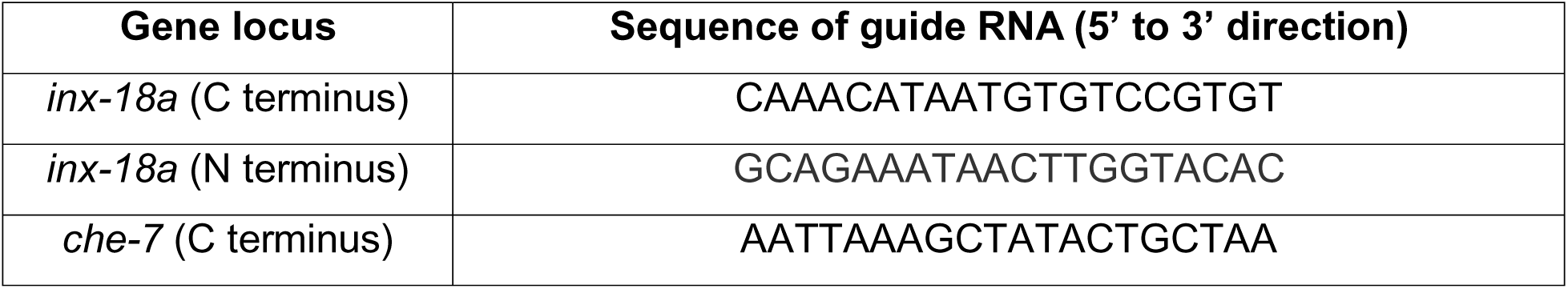

### Generation of Plasmids/ Constructs

All the plasmids used in this study are listed in Table S2.

### Generation of GLR nuclear reporter construct

For the ease of making nuclear reporter constructs, GFP encoding sequence in pPD95.75 vector was replaced with a *his-24::mCherry::unc-54 3’UTR* cassette to generate pAS010. GLR-specific cis-regulatory sequence *nep-2p7* (150 bp) (Stefanakis et al. 2024), was amplified from N2 genomic DNA and cloned into the pAS010 upstream of *his-24::mCherry* using PstI and MscI restriction endonuclease digestion to generate the *nep-2p7::his-24::mCherry::unc-54 UTR*.

### Generation of Cre and SplitGFP(1-10) constructs

GLR-specific *nep-2p7* driver sequence was cloned into *mec-18p::3XNLS::Cre::t2a::bfp* and *mec-18p::splitGFP(1-10)::t2a::bfp* constructs (Vats et al. 2026), replacing *mec-18p* sequence using SphI and MscI restriction enzyme sites to generate *nep-2p7::3XNLS::Cre::t2a::bfp* and *nep-2p7::splitGFP(1-10)::t2a::bfp.* To generate RME-specific *splitGFP(1-10)::t2a::bfp* expression constructs *unc-25p* (225 bp) cis-regulatory sequence^59^ was amplified from the genomic DNA and cloned similarly.

### Generation of GLR-specific rescue constructs

ORF of the a-isoform of *inx-18* was amplified from N2 using the SuperScript III First-Strand Synthesis kit (Invitrogen, cat. No. 18080051) and gene-specific primers. ORFs were cloned downstream of the *nep-2p7* driver using the Gibson assembly protocol (NEB). For C-terminally *gfp-*tagged *inx-18a, gfp* coding sequence was cloned before the STOP codon.

The plasmid expressing (pAN62) *unc-7-mPANX1*-*ECL2*^44^ was a gift from Dr. Shunji Nakano. *unc-7-mPANX1*-*ECL2* cassette from this plasmid was cloned downstream of the *nep-2p7* driver in the pPD95.75 vector backbone to generate GLR-specific expression construct: *nep-2p7::unc-7-mPANX1*-*ECL2*. A sequence encoding GFP was cloned downstream of the *unc-7-mPANX1*-*ECL2* cassette before the stop codon using a PCR fusion technique^60^ to generate the *nep-2p7::unc-7-mPANX1-ECL2::gfp* construct.

To generate the chimeric *inx-18a-mPANX1-ECL2*, the structure of INX-18 was predicted using DeepTMHMM 1.0 - BIOINFORMATIC SERVICES (Hallgren et al. 2022), and the *inx-18* sequence corresponding to the 2^nd^ extracellular loop (amino acids 221 to 287) was replaced with mPANX1 EC-loop 2 as previously published^44^ (Nakayama et al. 2024). PCR fusion technique^60^ was used to generate GLR-specific *nep-2p7::inx-18a-mPANX1-ECL2* construct*. nep-2p7::inx-18a-mPANX1-ECL2::gfp* construct was derived from the above construct by cloning the *gfp* encoding sequence before the stop codon, using PCR fusion technique^60^.

### Generation of GLR-specific Calbindin construct

A gene fragment containing the human Calbindin (CALB D-28K) sequence [based on Origene Technologies, Inc. (Accession Number: NM_004929)] (Schumacher et al. 2012), which is codon optimized for *C. elegans,* was obtained from Integrated DNA Technologies (IDT). This fragment was cloned downstream of the *nep-2p7* driver, followed by a t2a::*ebfp::unc-54 3’ UTR* cassette in the pPD95.75 vector backbone using KpnI and NcoI Restriction enzyme sites.

### BAPTA-AM Treatment

5g of BAPTA-AM (Millipore, Catalog no. 196419) was dissolved in 50µL of 100% DMSO (Sigma-Aldrich, Catalog no. 34869) to make a 131mM BAPTA-AM stock solution.

Synchronized populations of well-fed L4 stage animals, maintained at 20°C, were washed and resuspended in 1mL M9 buffer containing 7.6µL of BAPTA-AM stock solution and pelleted *E. coli* OP50 from 2mL of overnight culture (Resultant composition: 1mM final concentration of BAPTA-AM in M9 and *E. coli* OP50 pellet from 2mL culture). As a control, animals were treated with an equivalent volume of DMSO, i.e., 1mL of 0.76% v/v DMSO and *E. coli* OP50 pellet from a 2mL culture). The animals were incubated for 12 hours at room temperature (22°C) on a nutator. Treated animals were washed with M9 and imaged immediately.

### Salt Hypersensitivity Assays

All assays were performed with 1-day-old adult animals at 20°C, as previously described by (Stefanakis et al. 2024). To avoid potential variability, all strains in a comparison, including wild type, were recorded simultaneously. For each assay ∼20 animals were placed with eyelash pick on freshly made unseeded NGM plates containing 200mM NaCl. *floxed-unc-7* and *tdc-1(n3240)* mutant animals along with wild type controls were also tested in presence of 225 mM NaCl and were found to behave similarly. Animals were scored for their recovery from salt-induced paralysis within a time span of 60 minutes. The videos were captured using WormTracker (MBF Bioscience) image capturing software and data points were plotted and analyzed using GraphPad Prism 8.0.

### Microscopy, Image Processing and Data Analysis

Animals were anesthetized with 100 mM Sodium Azide and mounted on 5% agarose pads on glass slides. Images were taken using Olympus FV3000 or Nikon AX point-scanning confocal microscopes. Image processing and analysis were done by scanning the full Z-stack using NIH Fiji software. Maximum intensity projections of relevant z sections for all representative images and for GLR length quantification were done using the NIH Fiji software. Figures were prepared using Adobe Photoshop 2025 and Adobe Illustrator 2025. Separate channels were usually adjusted independently using Levels and Curves in Adobe Photoshop.

For length analysis of GLR cells, the lengths of all individual GLRs per animal were traced and measured using the segmented line tool in Fiji. The average length of all projections per animal was recorded and normalized to the distance from the grinder to the nose tip of the same animal. To assess GLR glial cell length in the presence of extrachromosomal rescue transgenes or Calbindin overexpression constructs, a *nep-2p7::ebfp2* construct was co-injected, and only cells expressing BFP were quantified and averaged per animal. Animals were stage-matched as 1-day-old adults for morphological assays.

### Quantification and Statistical Analysis

Statistical tests were applied based on the type of data set, distribution, and sample size. Groups were tested for normal distribution using the Shapiro-Wilk test. For comparisons between two groups following non-normal distributions, the Kolmogorov Smirnov-test was used, while in cases of normal distribution, unpaired t-test with Welch’s correction was used. For salt-hypersensitivity assays, multiple t-tests followed by a two-stage step-up procedure of Benjamini, Krieger, and Yekutieli correction, with FDR(Q) = 5%, was applied to correct for false discovery rates. Pairwise Fisher’s exact test was applied for categorical data like SNB-1 defects in RME neurons. In all tests, p<0.05 was considered statistically significant.

## ACKNOWLEDGMENTS

We thank Selvanayaki Eswaramoorthy for *C. elegans* injections; Mrunmayee Lele and A Josey Lourdes for their help with molecular biology and behavioural assays; we thank wormbase.org and wormwiring.org for resources, and Caenorhabditis Genetics Center (CGC) (NIH P40OD010440), Kavita Babu, Shunji Nakano, and Kota Mizumoto for *C. elegans* strains and constructs; Kavita Babu and Urmi Bandyopadyay for comments on manuscript. We thank Central Imaging and Flow Cytometry, Sequencing, and *C. elegans* facilities at NCBS. This work was supported by DBT Wellcome Trust India Alliance (IA/I/20/2/505211) and by the Department of Atomic Energy, Government of India (Project Identification No. RTI-4018).

## AUTHOR CONTRIBUTIONS

A.S. and A.B. designed experiments and oversaw the project. A.S., and M.X.M. performed experiments. A.S., M.X.M., A. Bandyopadhyay, and S.R. contributed to *C. elegans* strain generations; A.B. obtained funding; A.S. and A.B. analyzed and interpreted data; A.S. and A.B. wrote the paper.

## SUPPLEMENTARY FIGURE LEGENDS

**Figure S1: UNC-1, but not CHE-7, localizes in a punctate pattern within GLR cells**

(A) Schematics showing the 5Xsplit-GFP-based *che-*7 and *unc-1* endogenous reporter alleles*, che-7(amz33[che-7::5xgfp11])*, and *unc-1(amz22[unc-1::5xgfp11]),* respectively.

(B) Punctate localization of endogenously split-GFP-tagged UNC-1 (yellow arrowheads) within the leaf-like projections of lateral GLR glial cells, marked by BFP expression (*nep2p7::gfp1-10::t2a::ebfp2*), pseudocoloured in magenta.

(C) Endogenously split-GFP-tagged CHE-7 failed to show detectable expression within the leaf-like projections of GLR glial cells (yellow dotted circle), marked by BFP expression (*nep2p7::gfp1-10::t2a::ebfp2*), (pseudocoloured in magenta).

(D) Pan-neuronal expression of GFP1-10 (*UPNp::gfp1-10::t2a::ebfp2*) showed punctate localization of endogenously split-GFP-tagged CHE-7 (yellow arrowheads) within the nerve ring. Neurons are marked by BFP expression (pseudocoloured in magenta).

**Figure S2: *che-7* does not affect salt-hypersensitivity behaviour or GLR glial length**

(A, C and D) Statistical comparisons were performed using multiple t-tests followed by two-stage step-up procedure of Benjamini, Krieger and Yekutieli correction, with FDR(Q)=5%. Distributions shown are standard error of mean.*p<0.05, ***p<0.001,–compared to wild-type animals.

(A) Mutation in *che-(ok2373)* did not affect paralysis or post-paralysis recovery of motility when animals were exposed to 200 mM NaCl, compared with wild-type animals.

(B) Schematic showing the floxed allele of *inx-18a(amz71amz78[loxP::inx-18a::loxP])*The brown line indicates the region deleted in *inx-18(ok2454)*.

(C) Animals with GLR-specific expression of Cre recombinase in *nep-2p7::NLS::Cre::t2a::ebfp2* exhibit slightly less paralysis and unaffected post-paralysis recovery of motility when exposed to 200 mM NaCl, compared with wild-type animals. Animals homozygous for the floxed allele in *inx-18* show unaffected paralysis or post-paralysis recovery when exposed to 200 mM NaCl, compared with wild-type animals.

(D) Cre recombinase-mediated, GLR-specific deletion of *unc-7* in loxP-flanked *unc-7* animals, *unc-7(amz14ot895[loxP::unc-7::tagRFP-t::loxP]),* had no significant effect on post-paralysis recovery of motility when exposed to 225 mM NaCl.

(E) Each point in the violin plot indicates the mean length of GLRs within a single animal, normalized to the distance between the nose tip of the animal and the grinder in the posterior pharynx within the same animal. The *che(ok2373)* mutation did not significantly affect the GLR glia length. Red horizontal line denote the median, and grey lines indicate quartiles. Statistical comparisons between groups were performed using unpaired t-test with Welch’s correction. *p<0.05. n=number of animals, specified beneath each distribution.

(F) Each point in the violin plot represents the distance from the grinder in the posterior pharynx to the RME neurite, normalized to the distance between the centre of anterior pharyngeal bulb and the grinder in the posterior pharynx within the same animal. The *unc-7(e5)* mutation, but not the *inx-18(ok2454)*, significantly altered the RME neurite position. Black horizontal lines denote the median, and grey lines indicate quartiles. Statistical comparisons between groups were performed using unpaired t-test with Welch’s correction test. n=number of animals, specified beneath each distribution. ns=non-significant, ****p<0.0001 – compared to wild-type animals.

**Figure S3: INX-18 forms independent, undocked hemichannels within GLR glial cells**

(A, C, E) GLR-specific expression of GFP-tagged INX-18 in *nep-2p7::inx-18a::gfp* results in punctate localization of INX-18::GFP (yellow arrowheads) within the leaf-like projection in lateral glial cells (pseudocoloured in magenta) in wild-type animals. Punctate localization of INX-18::GFP remained unaffected in *inx-1(amz11)* and *inx-7(ok2319)* mutant animals.

(B, D, F) Violin plots show quantifications of the data shown in Panel A, C, E, respectively. Statistical comparisons between groups were performed using Kolmogorov-Smirnov test. ns=non-significant. n=number of glial cells, specified beneath each distribution.

(G) Probabilities of N- and C-terminal intracellular (magenta), 1 through 4 transmembrane (red), 1st and 2nd extracellular loops (blue), and 3rd intracellular loop (magenta) domains within INX-18a, according to the DeepTMHMM prediction tool. Numbers indicate the corresponding amino acid positions.

(H) GLR-specific expression of chimeric UNC-7-PANX1-ECL2 in *unc-7(e5)* mutant animals failed to rescue the delayed recovery of motility after paralysis when exposed to 200 mM NaCl, as observed in these mutants. Statistical comparisons between groups at each time point were performed using multiple t-tests followed by two-stage step-up procedure of Benjamini, Krieger and Yekutieli correction, with FDR(Q)=5%. Distributions shown are standard error of mean. ns=non-significant, *p<0.05, **p<0.01, ***p<0.001, ****p<0.0001 – compared to wild-type animals.

**Figure S4: Salt hypersensitivity behaviour is independent of CDK-5 and PDE-4 activity.**

(A,C) Statistical comparisons were performed using multiple t-tests followed by two-stage step-up procedure of Benjamini, Krieger and Yekutieli correction, with FDR(Q)=5%. Distributions shown are standard error of mean.

(A) Mutations in *cdk-5(ok626)* do not affect paralysis or post-paralysis recovery of motility when exposed to 200 mM NaCl, compared with wild-type animals.

(B) Each point in the violin plot indicates the normalized average length of GLRs within a single animal. Animals mutant for *pde-4(ce628)* had no significant defect in GLR length. GLR lengths were normalized to the distance between the nose tip of the animal and the grinder in the posterior pharynx. Red horizontal line denotes the median, and grey lines indicate quartiles. Statistical comparisons between groups were performed using unpaired t-test with Welch’s correction. n=number of animals, specified beneath each distribution. ns=non-significant.

(C) Mutations in *pde-4(ce268)* do not affect paralysis or post-paralysis recovery of motility when exposed to 200 mM NaCl, compared with wild-type animals.

