## Supplementary figures and images for "Molecularly Distinct Innexin Gap Junction Channels and Undocked Hemichannels Regulate Glia Morphology and Function in *Caenorhabditis elegans*"

### Supplemental Figures S1 - S4

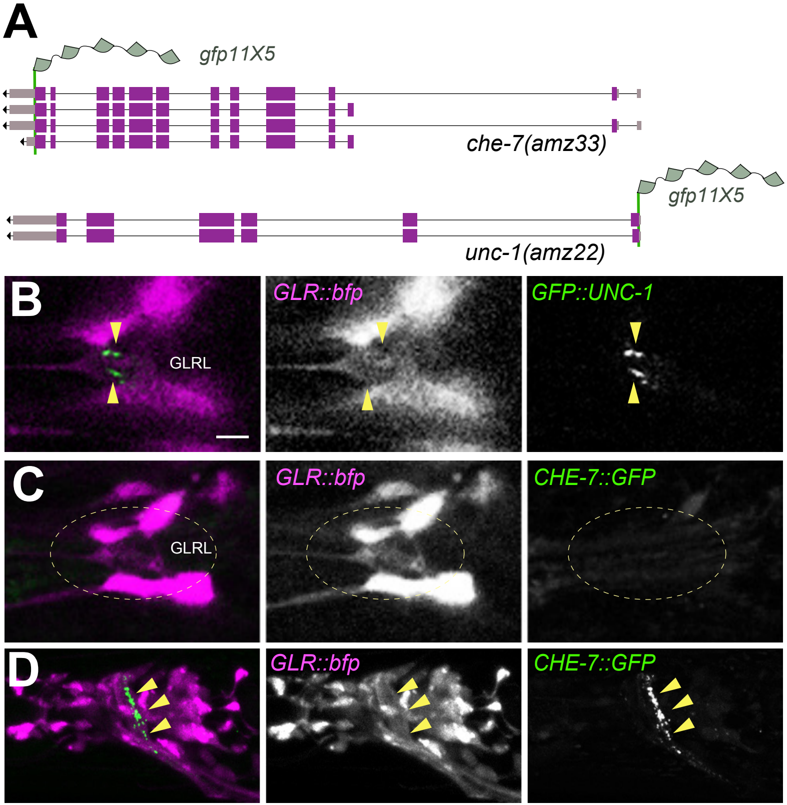

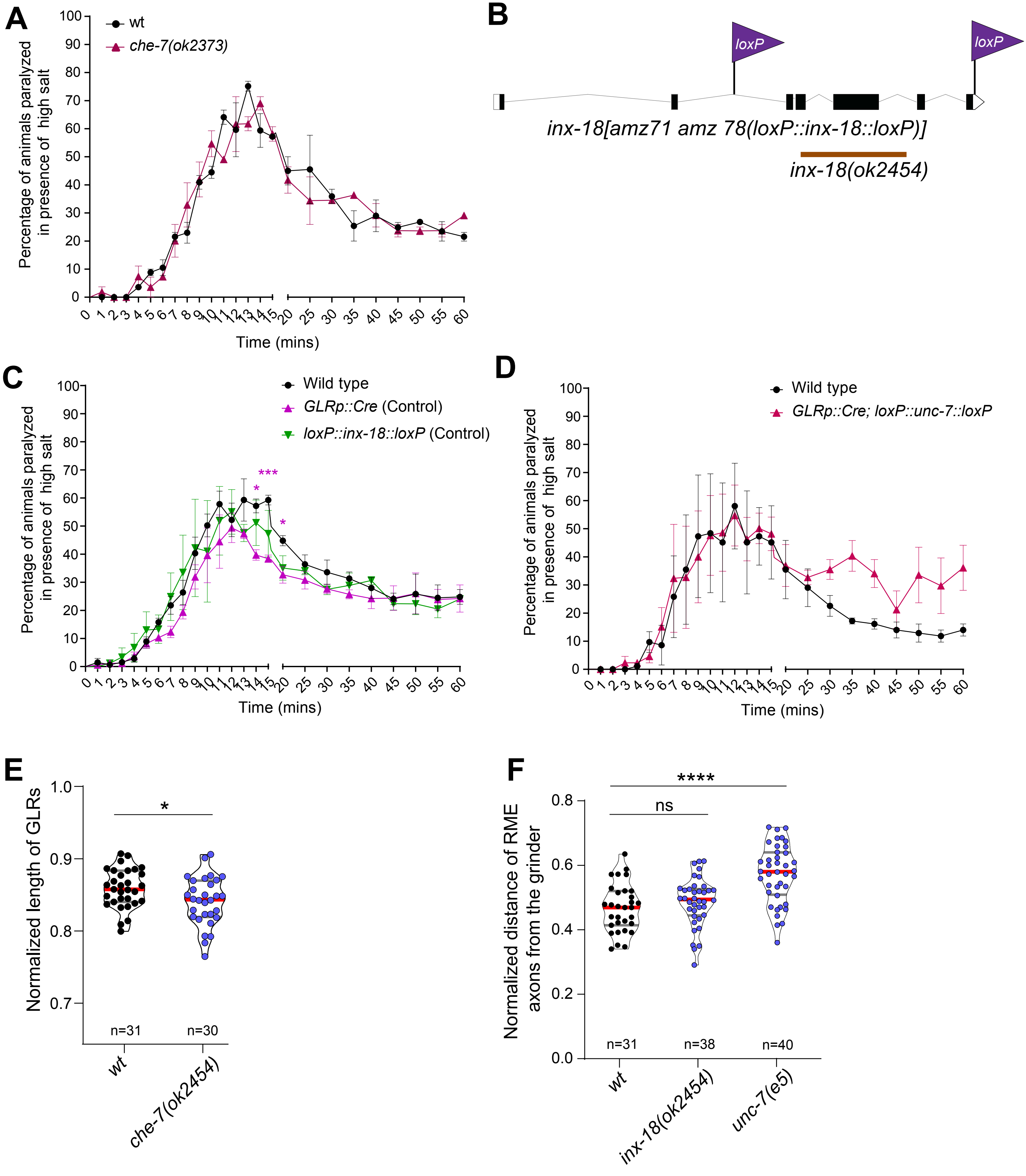

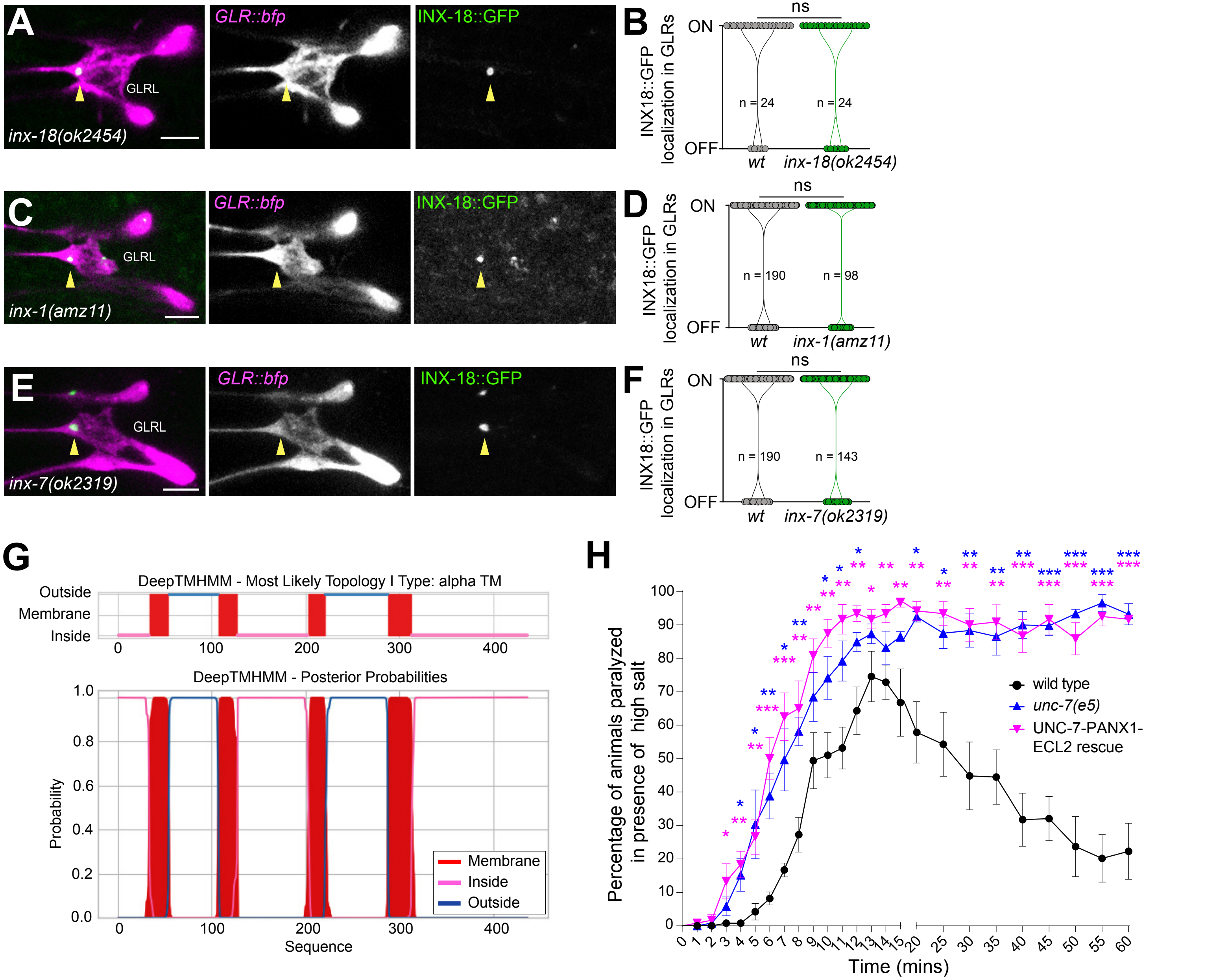

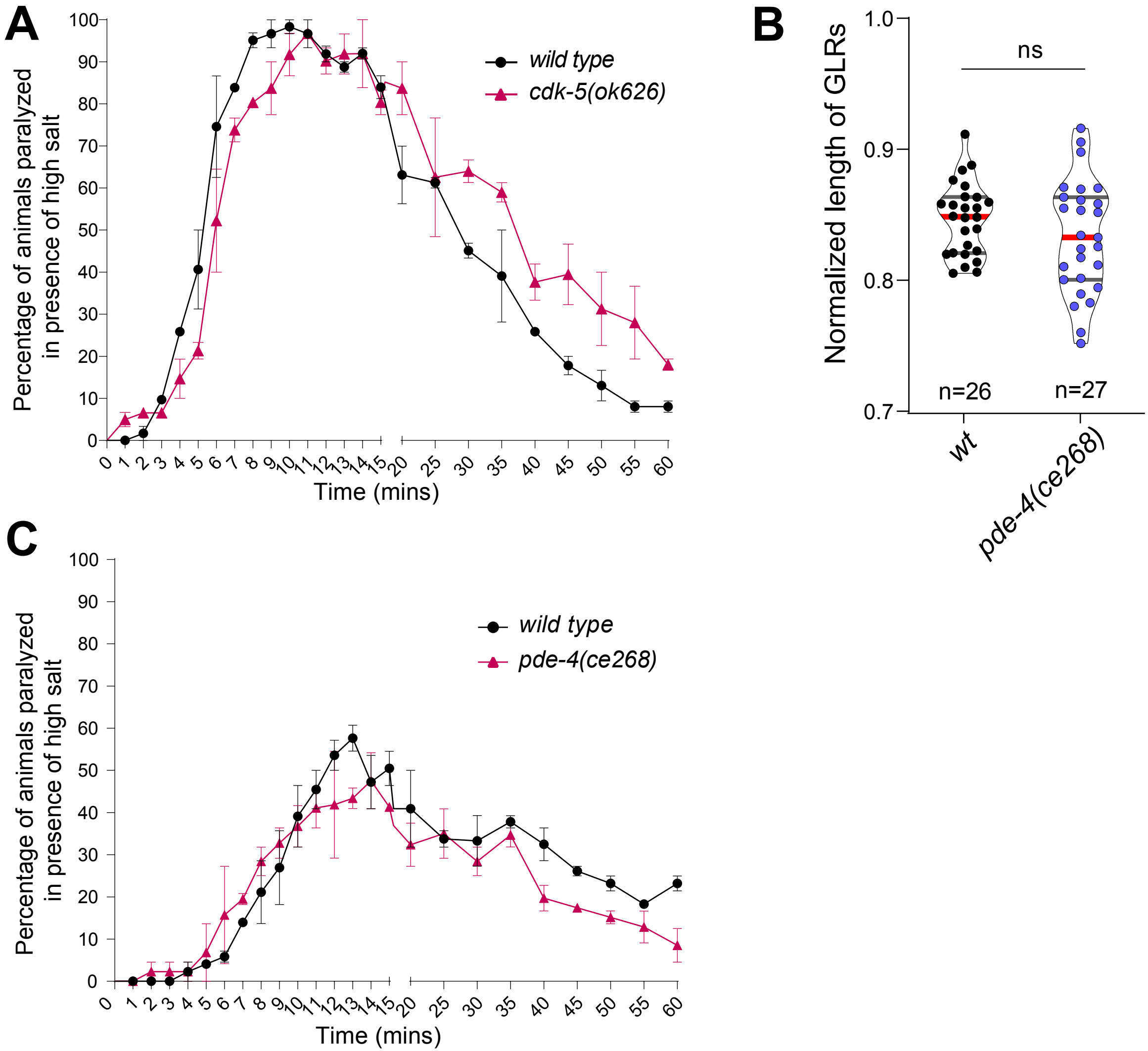
