## Supplementary material for "Molecularly Distinct Innexin Gap Junction Channels and Undocked Hemichannels Regulate Glia Morphology and Function in *Caenorhabditis elegans*": Table S1

| Sl. No. | Strain Name | Strain Genotype | Source |
|---|---|---|---|
| 1 | N2 | *C. elegans* wild isolate (variety *Bristol*) | CGC |
| 2 | ABH201 | *amzIs11[nep-2p7::his-24::mCherry; unc-122::GFP]* | This study |
| 3 | ABH232 | *amzIs12[nep-2p7::his-24::mCherry; unc-122::GFP]; otIs771(otEx7075[inx-18a WRM0629cH03fosmid::SL2::NLS::yfp::H2B)]); pha-1* | This study |
| 4 | ABH294 | *amzIs12[nep-2p7::his-24::mCherry; unc-122::GFP], che-7(syb4693[che-7::SL2::GFP::H2B]) V* | This study |
| 5 | ABH230 | *amzIs12[nep-2p7::his-24::mCherry; unc-122::GFP]; inx-7(ot904 [inx-7::sl2::gfp::h2b])* | This study |
| 6 | ABH233 | *amzIs12[nep-2p7::his-24::mCherry; unc-122::GFP]; inx-13(syb6884[inx-13::sl2::gfp::h2b])* | This study |
| 7 | ABH234 | *amzIs12[nep-2p7::his-24::mCherry; unc-122::GFP]; otIs756[otEx7102 [inx-5_fosmid::sl2::yfp::h2b; pha-1(+); myo-2::bfp]; pha-1* | This study |
| 8 | ABH205 | *unc-7(amz15[unc-7::splitGFP(11x5)])* | *Vats et al., 2026*[1] |
| 9 | ABH388 | *inx-18a[amz61(inx-18a::sfGFP(11X5)] IV* | This study |
| 10 | UJ1300 | *unc-9(miz81[unc-9::splitGFP(11X7)_LoxP])* | *Hendi et al., 2022*[2] |
| 11 | ABH249 | *unc-1(amz22[sfGFP(11X5)::unc-1]) X* | *Vats et al., 2026*[1] |
| 12 | ABH284 | *che-7(amz33[che-7::splitGFP(11x5)])* | *Vats et al., 2026*[1] |
| 13 | ABH379 | *amzIs17[nep-2p7::splitGFP(1-10)::t2a::ceBFP2, unc-122::GFP], inx-18a[amz61(inx-18a::sfGFP(11X5))] IV* | This study |
| 14 | ABH405 | *amzIs17[nep-2p7::splitGFP(1-10)::t2a::BFP, unc-122::GFP], unc-7(amz15[unc-7::splitGFP(11x5)])* | This study |
| 15 | ABH406 | *amzIs17[nep-2p7::splitGFP(1-10)::t2a::ceBFP2, unc-122::GFP], unc-9(miz81[unc-9::splitGFP(11X7)_LoxP])* | This study |
| 16 | ABH407 | *amzIs17[nep-2p7::splitGFP(1-10)::t2a::ceBFP2, unc-122::GFP], unc-1(amz22[sfGFP(11X5)::unc-1]) X* | This study |
| 17 | ABH456 | *amzIs17[nep-2p7::splitGFP(1-10)::t2a::BFP, unc-122::GFP], che-7(amz33[che-7::splitGFP(11x5)])* | This study |
| 18 | OH17075 | *inx-18(ok2454)* | *CGC* |
| 19 | OH13924 | *che-7(ok2373) V.* | Bhattacharya *et al., 2019*[3] |
| 20 | CB5 | *e5* | CGC |
| 21 | CB101 | *e101* | CGC |
| 22 | ABH495 | *amzIs19(nep-2p7::inx-18 cDNA::GFP, unc-122::GFP)* | This study |
| 23 | ABH334 | *nsIs700 [nep-2(prom7)::tagRFP] V, inx-18(ok2454)* | This study |
| 24 | ABH487 | *nsIs700 [nep-2(prom7)::tagRFP]; unc-7(e5) X.* | This study |
| 25 | ABH428 | *nIs213[egl-6::gfp; lin-15AB(+)], che-7(ok2373) V* | This study |
| 26 | ABH459 | *nsIs700 [nep-2(prom7)::tagRFP] V, juIs1 [unc-25p::snb-1::GFP + lin-15(+)] IV, inx-18(ok2454) IV, unc-7(e5)* | This study |
| 27 | CZ333 | *juIs1 [unc-25p::snb-1::GFP + lin-15(+)] IV.* | CGC |
| 28 | ABH438 | *juIs1 [unc-25p::snb-1::GFP + lin-15(+)] IV, inx-18(ok2454) IV* | This study |
| 29 | ABH437 | *juIs1 [unc-25p::snb-1::GFP + lin-15(+)] IV, unc-7(e5) X* | This study, and *Meng et al 2016*[4] |
| 30 | ABH475 | *amzEx142[nep-2p7::Cre::t2a::BFP, ttx-3::gfp], inx-18a(amz71 amz78[5' loxP::inx-18a::loxP 3']) IV* | This study |
| 31 | ABH415 | *amzEx142[nep-2p7::Cre::t2a::BFP, ttx-3::gfp]* | This study |
| 32 | ABH511 | *amzEx142[nep-2p7::Cre::t2a::BFP, ttx-3::gfp]; unc-7[amz14 ot895[5' loxP::unc-7::tagRFP-t::loxP]* | This study |
| 33 | ABH237 | *unc-7[amz14 ot895[5' loxP::unc-7::tagRFP-t::loxP]* | *Vats et al., 2026*[1] |

| | | | |
|---|---|---|---|
| 34 | ABH492 | *amzEx154(nep-2p7::inx-18-mPANX1::UTR, nep-2p7::sfGFP(1-10)::t2a::BFP, unc-122::GFP),nsIs700 [nep-2(prom7)::tagRFP], inx-18(ok2454) (#2)* | This study |
| 35 | ABH514 | *AmzEx156(nep-2p7::inx-18-mPANX1::UTR, nep-2p7::sfGFP(1-10)::t2a::BFP, unc-122::GFP),nsIs700 [nep-2(prom7)::tagRFP], inx-18(ok2454) (#1)* | This study |
| 36 | ABH493 | *nsIs700 [nep-2(prom7)::tagRFP]; unc-7(e5) X; amzEx155[nep2p7::unc-7::PANX1::GFP; nep2p7::sfGFP(1-10)::t2a::BFP; unc-122p::GFP)]* | This study |
| 37 | ABH504 | *amzIs19(nep-2p7::inx-18 cDNA::GFP, unc-122::GFP), inx-18(ok2454), nsIs700 [nep-2(prom7)::tagRFP] V* | This study |
| 38 | ABH451 | *nsIs700 [nep-2(prom7)::tagRFP], amzIs17[nep-2p7::splitGFP(1-10)::t2a::BFP, unc-122::GFP], inx-18a[amz61(inx-18a::sfGFP(11X5))] IV, unc-7(e5)* | This study |
| 39 | ABH515 | *nsIs700 [nep-2(prom7)::tagRFP], amzIs17[nep-2p7::splitGFP(1-10)::t2a::BFP, unc-122::GFP], unc-7(amz15[unc-7::splitGFP(11x5)]), inx-18 (ok2454)* | This study |
| 40 | ABH513 | *amzIs19(nep-2p7::inx-18 cDNA::GFP, unc-122::GFP), unc-9(e101), nsIs700 [nep-2(prom7)::tagRFP] V* | This study |
| 41 | ABH509 | *amzIs19(nep-2p7::inx-18 cDNA::GFP, unc-122::GFP), inx-7(ok2319)* | This study |
| 42 | ABH510 | *amzIs19(nep-2p7::inx-18 cDNA::GFP, unc-122::GFP), inx-1(amz11)* | This study |
| 43 | ABH490 | *nsIs700 [nep-2(prom7)::tagRFP]; unc-7(e5) X; amzEx152[nep2p7::unc-7::PANX1; nep2p7::sfGFP(1-10)::t2a::BFP; unc-122p::GFP)] (#1)* | This study |
| 44 | ABH491 | *nsIs700 [nep-2(prom7)::tagRFP]; unc-7(e5) X; amzEx153[nep2p7::unc-7::PANX1; nep2p7::sfGFP(1-10)::t2a::BFP; unc-122p::GFP)] (#2)* | This study |
| 45 | ABH481 | *nsIs700 [nep-2(prom7)::tagRFP]; amzEx149[nep-2p7::Calbindin::t2a::BFP; nep-2p7::splitGFP(1-10)::t2a::BFP, unc-122p::GFP]* | This study |
| 46 | ABH464 | *nsIs700 [nep-2(prom7)::tagRFP] V, cdk-5(ok626) III* | This study |
| 47 | ABH463 | *nsIs700 [nep-2(prom7)::tagRFP] V, Y39A3CL.5(ok2808) III* | This study |
| 48 | RB2123 | *Y39A3CL.5(ok2808) III.* | CGC |
| 49 | RB814 | *cdk-5(ok626) III.* | CGC |
| 50 | ABH457 | *nsIs700 [nep-2(prom7)::tagRFP] V, pde-4(ce268) II* | This study |
| 51 | KG744 | *pde-4(ce268) II* | CGC |
| 52 | MT10661 | *tdc-1(n3240) II.* | CGC |
| 53 | MT16684 | *nIs213[egl-6::gfp; lin-15AB(+)]* | Ringstad et al., 2008[5] |
| 54 | OH15242 | *otEx7090 [inx-11(fosmid WRM0621cA09)::SL2::NLS::YFP::H2B + pha-1(+) + unc-122::GFP]* | Bhattacharya *et al., 2019*[3] |
| 55 | OH15282 | *otEx7113 [inx-12(fosmid WRM0621dC07)::SL2::NLS::YFP::H2B + pha-1(+) + myo-2p::BFP]* | Bhattacharya *et al., 2019*[3] |
| 56 | OH15285 | *otEx7116 [inx-1A_fosmid::sl2::yfp::h2b; pha-1(+); myo-2::bfp]* | Bhattacharya *et al., 2019*[3] |
| 57 | OH15288 | *otEx7119 [inx-10A_fosmid::sl2::yfp::h2b; pha-1(+); myo-2::bfp]* | Bhattacharya *et al., 2019*[3] |
| 58 | OH15532 | *otEx7227[inx-9_fosmid::SL2::1xNL2::YFP::h2b; myo-2::BFP; pha-1(+)]* | Bhattacharya *et al., 2019*[3] |
| 59 | OH15546 | *inx-7(ot904 [inx-7::sl2::gfp::h2b])* | Bhattacharya *et al., 2019*[3] |

| | | | |
|---|---|---|---|
| 60 | OH15566 | *inx-2(ot906 [inx-2::sl2::yfp::h2b])* | Bhattacharya *et al., 2019*[3] |
| 61 | OH16524 | *otIs769(otEx7233[nsy-5_fosmid::sl2::yfp::h2b; myo-2::bfp; pha-1(+)])* | Bhattacharya *et al., 2019*[3] |
| 62 | OH16527 | *otIs772(otEx7106 [unc-7_fosmid::sl2::yfp::h2b; pha-1(+); myo-2::bfp])* | This study |
| 63 | OS11484 | *nsIs700 [nep-2(prom7)::tagRFP]* | *Stefanakis et al., 2024*[6] |
| 64 | PHX4693 | *che-7(syb4693[che-7::SL2::GFP::H2B]) V* | *CGC* |
