## Supplementary material for "Molecularly Distinct Innexin Gap Junction Channels and Undocked Hemichannels Regulate Glia Morphology and Function in *Caenorhabditis elegans*": Table S2

| Sl. No. | Plasmids used in this study |
|---|---|
| 1 | *MCS::5' fire vector intron::his-24::mCherry::unc-54 3' UTR* |
| 2 | *nep-2p7::5' fire vector intron::his-24::mCherry::unc-54 3' UTR* |
| 3 | *nep-2p7::splitGFP(1-10)::t2a::ceBFP2::p10 3' UTR* |
| 4 | *UPNp::splitGFP(1-10)::t2a::ceBFP2::p10 3' UTR* |
| 5 | *nep-2p7::inx-18(ORFs)::GFP::p10 3' UTR* |
| 6 | *nep-2p7::3xNLS::Cre::t2a::ceBFP2::p10 3' UTR* |
| 7 | *nep-2p7::inx-18-mPANX1-ECL2::p10 3' UTR* |
| 8 | *nep-2p7::unc-7-mPANX1-ECL2::unc-54 3' UTR* |
| 9 | *nep-2p7::Calbindin D28K::t2a::ceBFP2::p10 3' UTR* |
| 10 | *unc-122::GFP* |
| 11 | *ttx-3::GFP* |
